# Isolation of highly purified genomic material from mitochondria of muscle tissue cells

**DOI:** 10.1101/2022.12.12.520064

**Authors:** M.A. Tatarkina, V.V. Lobanova, I.I. I.I. Kozenkov, B.E. Efimenko, A. K. Dzhigkaev, K.Y. Popadin, K.V. Gunbin, A.G. Goncharov

## Abstract

In this work, we adapted a method for isolation of a highly purified fraction of mitochondrial DNA from muscle tissues suitable for further sample preparation of libraries without an amplification step for sequencing tasks on various NGS platforms and a method for evaluating the purity of DNA from contamination by nuclear genome regions.

We optimized several techniques^7,8^ for enrichment of the mitochondrial fractions and purifying mtDNA. Here, we describe a protocol that allows getting from 80-100 mg of muscle tissues up to 1000 ng mtDNA, almost free from impurities of RNA and fragments of the nuclear genome.To assess the degree of purity of human mtDNA fraction from impurities of the nuclear genome, we adapted the PCR-screening technique^7^ for the beta-actin gene region and AluSx-repeats in the human genome.

This methodology avoids false-heteroplasmy calls (PCR biases or NUMT contamination) that occur when long-range PCR amplification is used for mtDNA enrichment.

## INTRODUCTION

Aging is a highly complex process associated with numerous phenotypes. Somatic mutations in the mitochondrial genome (mtDNA) are considered one of the most critical aging factors due to the high rate of mtDNA mutagenesis and the essentiality of mitochondrial metabolism in all tissues. It has been shown, for example, that deleterious somatic mtDNA deletions in neurons and skeletal muscles are accumulated with age leading to neurodegeneration and muscular dystrophy. Non-proliferating (postmitotic) tissues are the most vulnerable to somatic mtDNA mutations due to the possibility of clonal expansion of selfish (with deleterious variant) mtDNA within a cell.

Here, to investigate deeper the mitochondrial component of aging, we established a biobank of the skeletal muscles of aged people. We collected and annotated hundreds of samples, performed PCR-free enrichment of mtDNA and sequenced the libraries deeply. As expected, we observed a positive relationship between the mtDNA mutational burden and age. Next, we focus on a high level of variation in mtDNA heteroplasmy, which is unexplained by age. Harmful mutations in the mitochondrial genome can be compensated for by an increased copy number of the mitochondrial genome ^1^

Population aging, namely the steady increase in the proportion of people over 65, is one of the most important social problems of the modern world. It is important to note that every year the imbalance in the proportions of age groups will only increase (WHO). Demographic aging raises questions for the scientific community about the mechanisms of aging development, prevention of diseases of the elderly and measures to prolong the period of “healthy” (active) aging. Among the numerous hypotheses and theories of aging, the so-called “mitochondrial theories of aging” occupy a special place since the development of senile changes is to some extent associated with a violation of the main function of mitochondria, namely the oxidation of nutrients and the production of ATP molecules - a universal source of energy for most biological processes in cells^2^.

The accumulation of aberrations in the mitochondrial genome with age seems to play a decisive role in intracellular metabolism disorders. At the same time, a high mutation rate ultimately leads to the accumulation of mitochondrial heteroplasmy in older people with age, which, in turn, disrupts the processes of mitophagy and, ultimately, leads to the development of immune-inflammatory reactions that underlie the pathogenesis of aging^3,4^

The possibilities of selecting determinants of mitochondrial heteroplasmy for subsequent analysis of its associations with age-related changes have significantly increased with the introduction of whole genome sequencing into routine scientific practice. Preparation of libraries for sequencing on NGS platforms requires a fraction of genomic material with a high degree of purification from concomitant contaminants - RNA, low molecular weight fraction of DNA, proteins or oligonucleotides. In the case of studying the genetic material of cell organelles (mitochondria, chloroplasts or plastids), additional purification is required from fragments of the nuclear genome, which often carries transposed segments of the organelle genome, such as, for example, NUMT (NUclear-encoded MiTochondrial segments - mtDNA nuclear-encoded regions).The method of preparing libraries without an amplification step (PCR-free) is increasingly being used to search for rare allele variants with a low frequency of occurrence. This method requires the preparation of large amounts of high-purity starting DNA, on the order of 200–1000 ng, but at the same time avoids amplification errors introduced by the polymerase enzyme.

The analysis of single nucleotide variations and indels in the mitochondrial genome in somatic tissues requires deep sequencing of the pure mitochondrial DNA^5^. Usually, the high quantity of target sequence for upstream NGS application, such as library preparation, is achieved by routine methods of enrichment, such as long-range PCR amplification that can introduce biases into heteroplasmy analysis^6^.

## METHODS

### Participants and tissue harboring

The initial material for studying was muscle tissue samples with a volume of 10 to 50 cubic millimeters obtained during a hip or knee arthroplasty operation. The collection of biopsy specimens of such a volume is safe, does not harm the health of the patient and does not affect the further process of recovery of the operated patient. The biopsy material was taken in the Department of Traumatology and Orthopedics of the Center for High Medical Technologies of the Ministry of Health of the Russian Federation (Kaliningrad). All studies were carried out in strict accordance with bioethical norms and rules, conclusions were obtained from the independent Ethical Committee of the IKBFU and the Ethics Committee of the Federal State Budgetary Institution “FTsVMT” of the Ministry of Health of the Russian Federation. In the operating room, observing the rules of asepsis, muscle samples were placed in test tubes containing sterile phosphate-buffered saline (PBS) with 0.05% sodium azide or a mixture of 1x [Amp+Kan+ChlAm], and within 3 hours they were delivered to the Genomic Research Center of the IKBFU. Next, samples were prepared for mtDNA isolation.

### Tissue lysis

Tissue samples were incubated in a solution with collagenase from crab pancreas (Biolot) at the rate of 0.75 units/mg of tissue for 24 hours. Collagenase breaks down collagen fibers in muscle tissue, allowing a suspension of free muscle cells to be obtained, which allows better tissue homogenization. A Dounce’s homogenizer was used with 30 strokes of pestle A (70 to 120 microns) and 20 strokes of pestle B (20 to 55 microns), all procedures were performed on ice.

### Mitochondria isolation and purification

Differential centrifugation, according to the protocol of M. Isocallio^9^, was performed to separate mitochondria from cellular debris and other organelles such as nuclei. The composition of the buffer includes sucrose and bovine serum albumin, which allows to separate mitochondria in the supernatant, the rest settles to the bottom.

### MtDNA extraction and purification

mtDNA extraction was performed according to B. Arbeithuber protocol^7^. At this stage, we used plasmid-safe ATP-dependent DNAse (Exonuclease V, Lucigene) and RNase A (Biolot). This DNAse cuts the chains of linear double-stranded DNA molecules or DNA-RNA duplexes without affecting the circular ones.

After the first treatment with DNase, the portion of nuclear DNA (nDNA) associated with histones remained in the sample. The addition of proteinase K makes it possible to destroy these complexes, and the second treatment with DNAse leads to cleavage of the isolated nDNA. Proteinase also cleaves fragments of mitochondrial membranes and intracellular protein structures. In addition, RNAse A was re-added to remove residues of RNA inside the mitochondria.

The reaction proceeds in a lysis buffer containing 1.45% NaCl and 1% SDS for 45-60 minutes, which ultimately leads to the destruction of mitochondria.

Isolation of mtDNA from the solution was carried out by the standard method using phenol-chloroform extraction, precipitation with 96% ethanol and washing with 70% ethanol.

### Verification of purity and quality of mtDNA

To assess the purity of the isolated mtDNA fraction from nuclear DNA contamination, we used PCR analysis for regions of the nuclear genome. Mitochondrial ND5 and TRNF region were the controls for mtDNA, and actb and AluSx were the controls for nDNA presence. For all PCR variants (relative to control and absolute measurements), we used the same primer pairs for the ND5, TRNF, act-b gene regions, and the AluSX region, respectively. Primer sequences were:

AluSx-F CTGTAATCCCAGCACTTTGG
AluSx-R CTCTGTCGCCTAGGCTGGAGTGCA
ND5_homo_F tgtgatatataaactcagacccaaa
ND5_homo_R tgggctattttctgctaggg
b_actinHF CACCATTGGCAATGAGCGGTTC b_actinHR AGGTCTTTGCGGATGTCCACGT
R(TRNF) tgtttatggggtgatgtgag
F(TRNF) gtttatgtagcttacctcctcaa
NGS library preparation

The library of mtDNA was prepared from 500 ng mitochondrial DNA with a KAPA HyperPlus Kit PRC-free (Roche), according to the manufacturer’s instructions. The protocol included the following steps: library preparation, hybridization, bead capture, washing, enrichment QC, sequencing, and pre- and post-capture multiplexing. Quantification analysis and assessment of the average size and length of the NGS libraries were performed using a Bioanalyzer assay (Agilent). Sequencing on the NGS libraries was performed by a NovaSeq (Illumina) paired-end 2 × 150-bp DNA sequencing platform with a NovaSeq 6000 S2 Reagent Kit v2 (300-cycle, Illumina), according to the manufacturer’s procedure.

## RESULTS

### 1. DNA samples

As a result of our work on adapting methods for enriching mitochondrial fractions and purifying mtDNA, it was possible to obtain up to 1000 ng of mtDNA free from RNA impurities and fragments of the nuclear genome from initial 80–100 mg of muscle tissues.

### 2. Concentrations and mtDNA copy numbers

We measured the relative copy number of mtDNA in the samples using real-time PCR with SYBR Green I intercalating dye (qPCRmix-HS SYBR-low ROX kit, Evrogen) against mtDNA controls that were examined for absolute copy number in ddPCR. As an internal reference sample, we used mtDNA samples from ddPCR experiment with known absolute copy number, titrated to initial concentrations of mtDNA at level 10ng/μl, 1ng/μl and 0.1ng/μl. We measured the absolute copy number of mtDNA and nDNA in the samples using the QIAcuity One digital PCR platform (Qiagen).

Amplification curves for real-time PCR, as well as primary data and data in tabular form, are available under the link in Supplements 1 (Suppl.1, Table 1 - link) and ddPCR data is available under [link_1].

Figure 1 shows the results of the reaction with tenfold dilutions (10ng, 1ng, 0.1ng of DNA per reaction). The non-optimized response efficiency with the PowerTrack™ SYBR Green Master Mix was 65.51%. The enrichment of the mtDNA fraction relative to the mtDNA in total DNA control samples averaged k=8.454 times. For some samples the enrichment was k=20 times.

**Figure 1.**
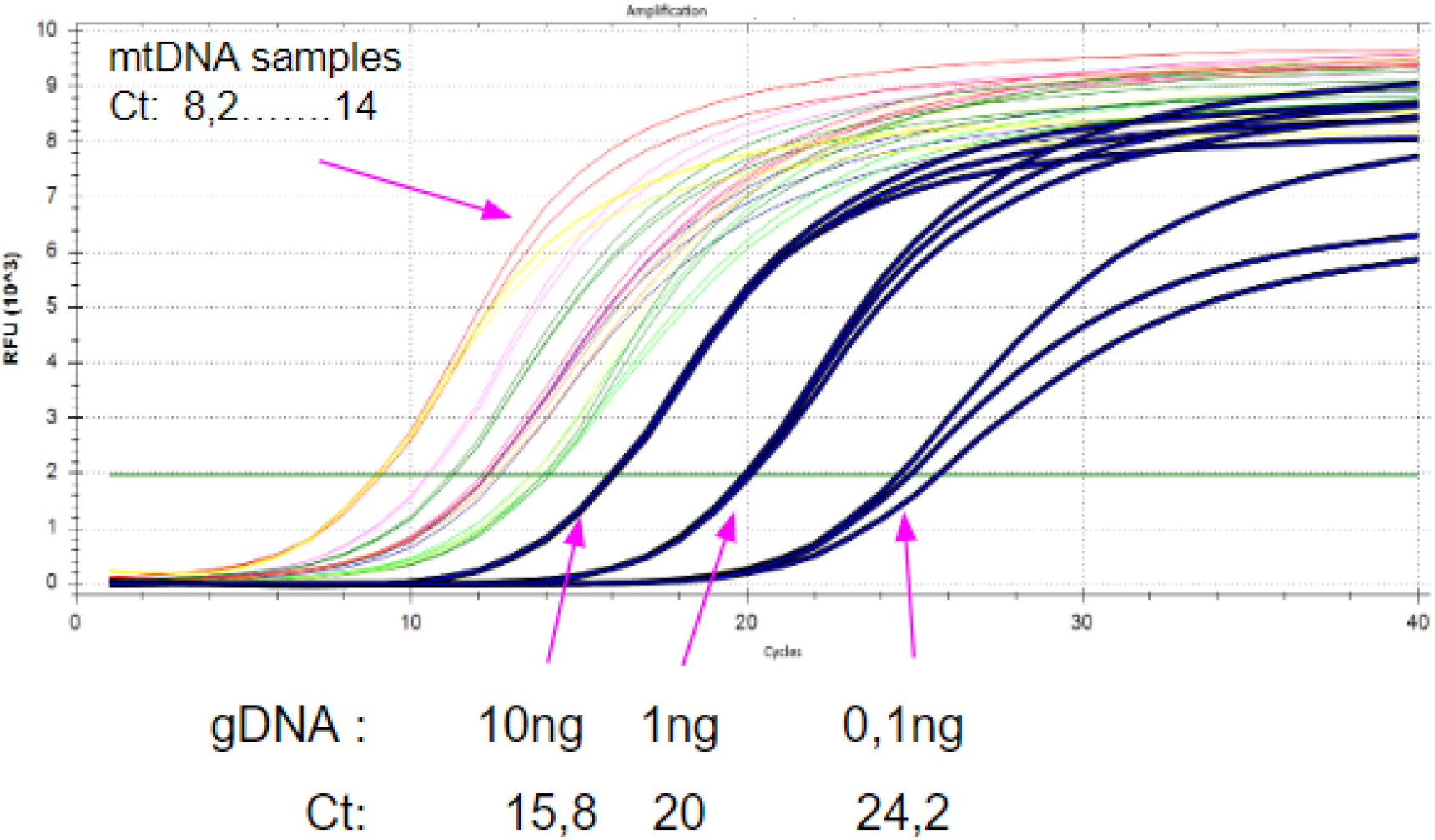
Amplification curves in real-time PCR.

Copy number was measured independently in ddPCR experiment and in sequence data analysis and results are presented at github (Suppl.1 GitHub_link_2)

### 3. mtDNA enrichment

To assess the degree of purity of the identified mtDNA from impurities of the large genome, we adapted the PCR screening method^2^ for the beta-actin gene region and Alu-repeats in the human genome.

HEK293 gDNA was used as the positive control, and no DNA was used as a template in the negative control. As presented in Figure 2 the weak bands 200-300 bp in size are visible in the lower part, thus we observe the contamination of the samples with gDNA material since in the track with the positive control (PC), a band of the amplified Alu-repeat region of similar brightness and size is visible. Lane 92* presented amplicons of the ND5 gene region for the fraction of mtDNA in debris after differential centrifugation step. As can be seen in lane 92*, we lose some of the mtDNA during isolation. 5 ukl of the PC was loaded into the lane and 15 ukl of samples was loaded in all other lanes. For densitometry samples against the mass standart (ladder) quantity (mass) of all samples was normalized to their volumes.

**Figure 2.**
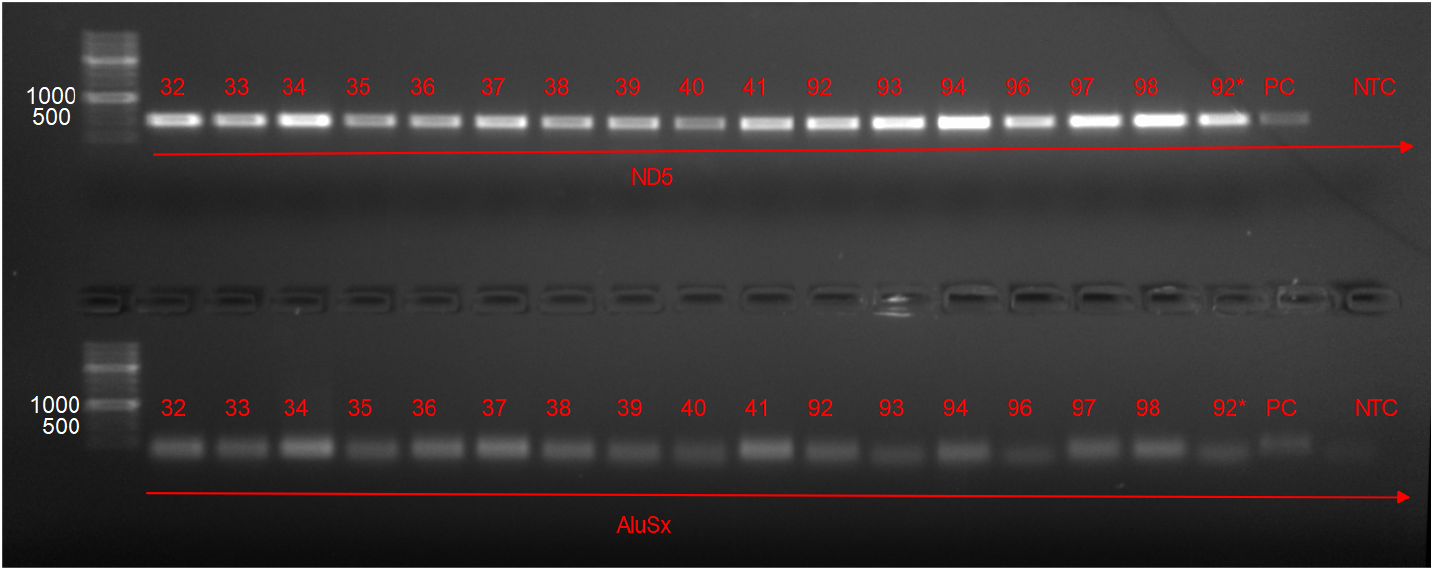
Results of gel electrophoresis for part of the samples, numbers on top. The upper part is amplification of the ND5 gene region, the lower part is amplification of Alu-repeat regions.

### 4. Preliminary sequencing results

First of all, 47 samples of sequencing data (pair-end reads) were filtered from low-quality reads and sectional reads using fastp 0.23.1 (options to run fastp: -g program --poly_g_min_len 5 -x --poly_x_min_len 10 −5 −3 -Z 4 -M 25 -n 3 -e 20 -l 77 -in -sh 12). Qualitative human observations were obtained for mapping to the reference mitochondrial genome (NC_012920.1). Mapping was carried out using the bwa 0.7.17-r1188 program. Using samtools 1.10 Received proportion of reads mapped to the mitochondrial genome, as well as genome coverage.

The proportion of mapped reads among sequenced samples ranges from 1.29% to 48.35% as shown in Figure 3.

**Figure 3.**
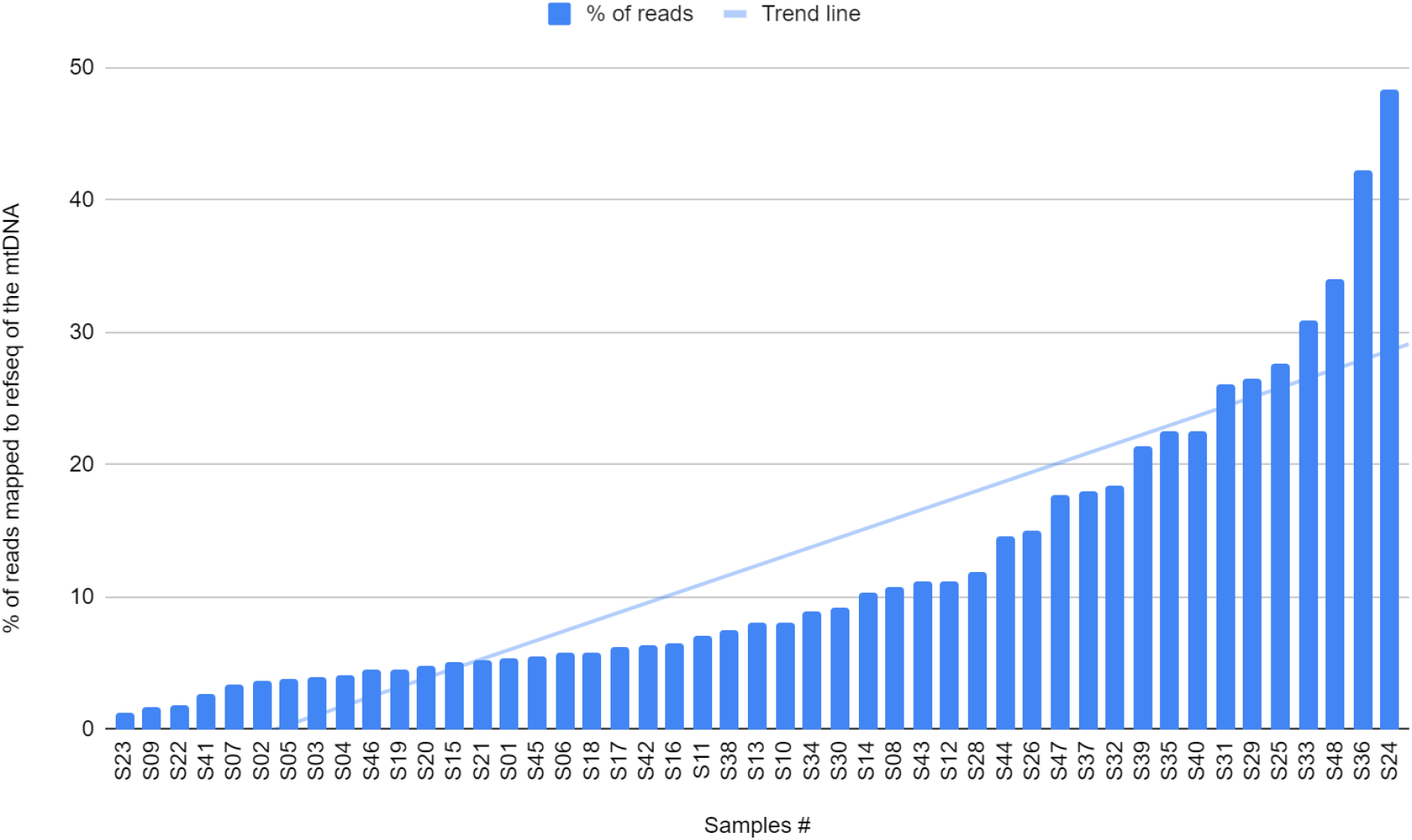
Percentage of reads mapped to the reference mitochondrial genome

Average genome coverage ranges from 12x to 3810x and for 34 samples (72%) is greater than 100x, 24 of which have coverage greater than 300x, data presented in Figure 4.

**Figure 4.**
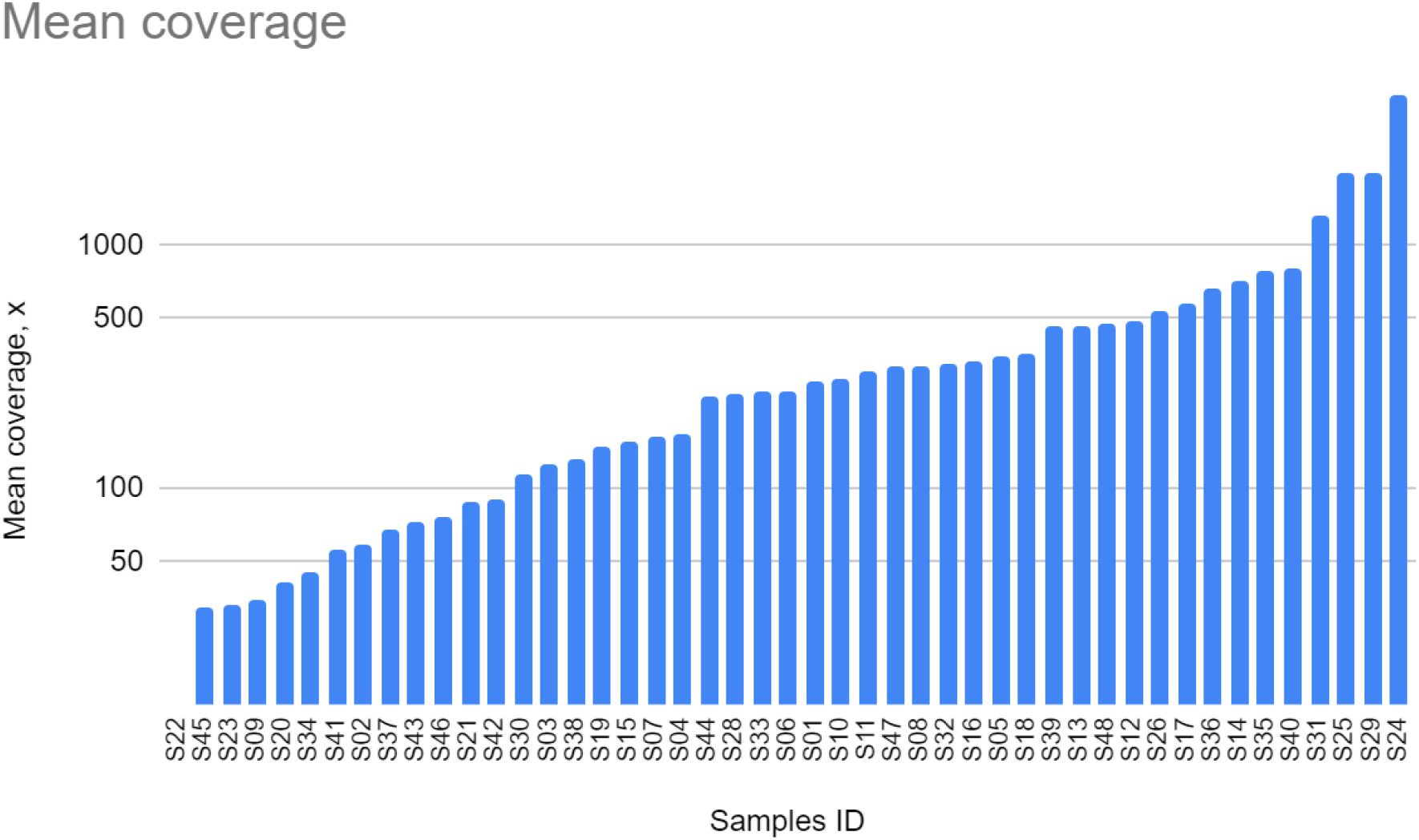
Mean coverage

## DISCUSSION

(A) A new hybrid protocol for the isolation of mitochondria from muscle tissue has been compiled. Several buffers and enzymes were tested for muscle tissue lysis, crab pancreas collagenase was chosen. A reproducible method for enrichment of the mtDNA fraction was worked out - with the isolation of intact organelles, subsequent double purification from impurities of linear nDNA and RNA, and lysis of the mitochondrial membrane.

Methods for controlling the purity of mtDNA relative to genomic DNA were tested - using PCR methods with amplification of AluSx repeat regions (introduced as a new reliable control), with controls for b-actin, ND5 gene regions.

(B) Differential centrifugation does not entirely eliminate nDNA and RNA from low-density suspension with intact mitochondria.

The original method used exonuclease V (ExoV/RecBCD) to eliminate genomic DNA. However, ExoV has exonuclease activity against both linear double-stranded or single-stranded DNA and against circular DNA molecules with a break in one of the strands. In this study gDNA contamination was detected with PCR amplification for AluSx regions - we settled on the AluSx repeat family and selected primers for the 5B and 3B repeat arms in order to amplify fragments from 200 to 600 bp. House-keeping genes such as 18S (18S rRNA), GAPDH, HPRT1, YWHAZ, UBC and RPII, POLR2A (RNA polymerase II) are commonly used for such analysis. But these genes are single-copy, i.e. in the entire nuclear DNA are found only once. In our work, we decided to use Alu-repeats as the target region - short dispersed nuclear elements (SINEs) from 100 to 900 bp in length, specific for primates. Approximately 10% of the human genome consists of Alu repeats^10^, which are fairly evenly distributed over all chromosomes. In the genomes of primates, the number of Alu repeats is more than 1 million copies^11^. That is why such repeats are a more reliable marker for nDNA detection by PCR than house-keeping genes.

AluSx - more relaible marker for gDNA contamination than other house-keeping genes usually presented in genome only in one or few copies.

Even if you physically removed a single chromosome (for example - chr7 with actb sequence as shown in Figure 5), you still can detect gDNA contamination in your sample with AluSx marker.

**Figure 5.**
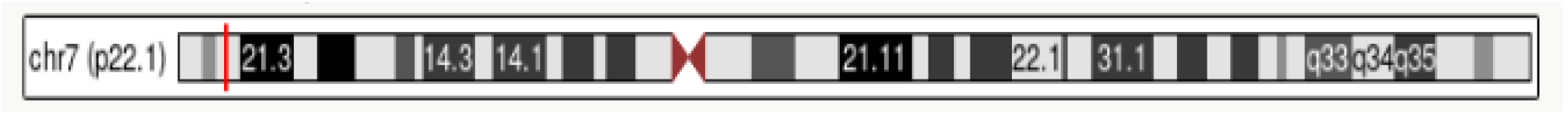
Physical site of actb sequence on chr7 marked with red line

(C) we achieved 8-20 fold mtDNA enrichment vs. gDNA, data shown in Figure 6.

**Figure 6.**
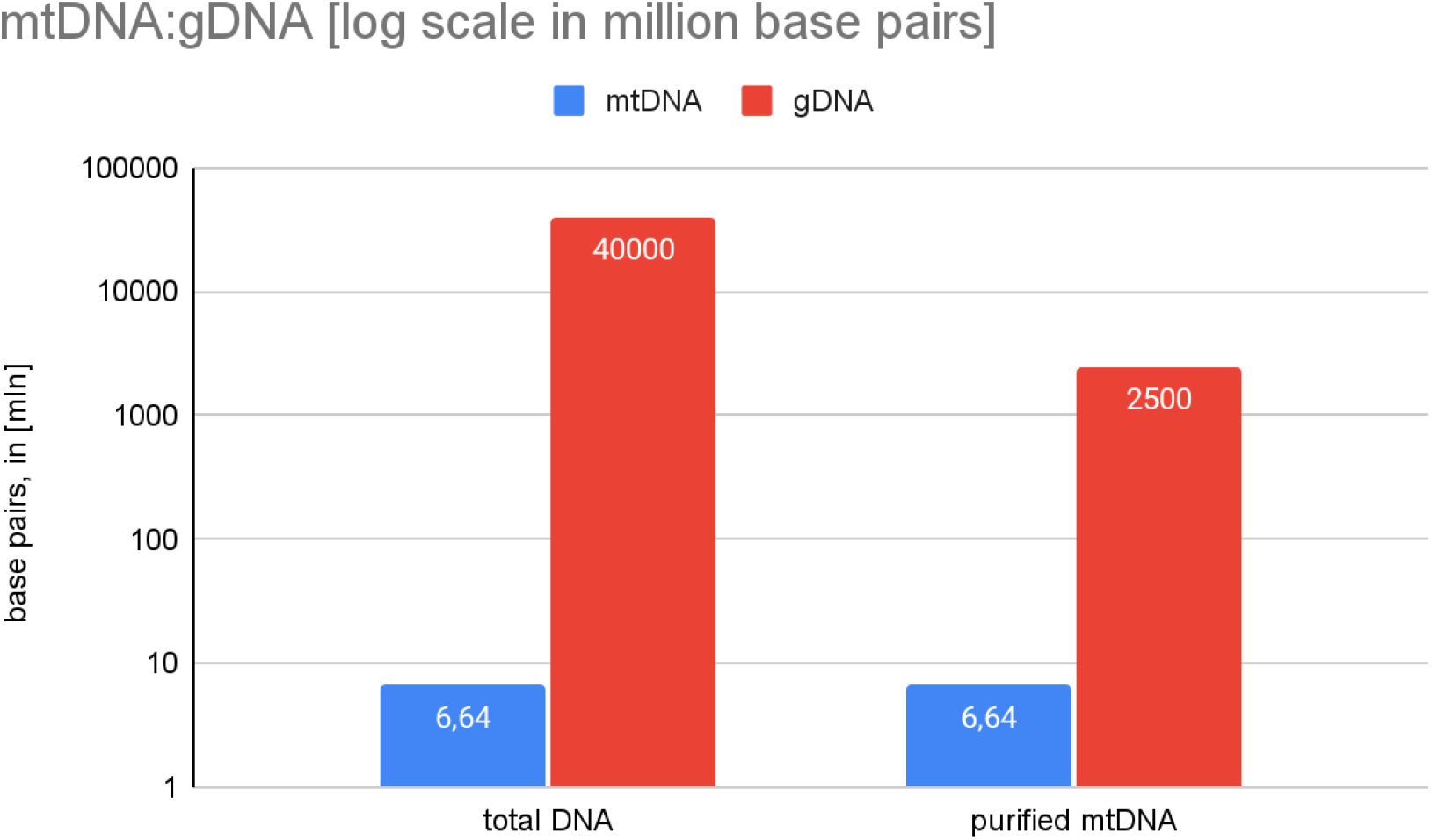
mtDNA enrichment against gDNA

In other words, if we initially have:

i. 400 copies mtDNA per cell (6,64 mln bp or ~0,01% of total bp in muscle cell) and
ii. 6-10 diploid nuclear genomes (muscle cells - multinuclear cells, 64 billion bp), then after enrichment we get in ideal case:
iii. 0,3 - 0,5 from diploid nuclear genomes (20-fold decrease, ~3,2 billion bp)
iv. the same 400 copies mtDNA (6,64 mln bp, but at this point = 0,2% of total bp in a sample from muscle cell).

We plan to uncover additional (to age) factors, potentially affecting the rate of mtDNA mutagenesis and clonal expansion of the somatic mtDNA mutations. Among such factors, we consider mtDNA haplogroups, nuclear genotypes as well as the lifestyle of participants. We propose that the somatic mutational burden of mtDNA can be a sensitive marker of biological age.

## ACKNOWLEDGMENTS

This work is supported by RSF No 21-75-20145.

## SUPPLEMENTARY FILES

Table 1. PCR amplification data (see additional file “LERA_2021-12-11 18-06-38_MTDNA_ANAL - Quantification Cq Results.xls”) - https://disk.yandex.ru/d/oWlOciWiOoxOpQ

link_1. ddPCR data https://disk.yandex.ru/d/9c9NnQEnziyo1Q

GitHub_link_2. Sequence data analysis and results https://github.com/mitoclub/OldMenGenetics/tree/master/data/processed/sea1_topmed

## Notes

https://github.com/mitoclub/OldMenGenetics/tree/master/data/processed/seq1_topmed

https://disk.yandex.ru/d/9c9NnQEnzjyo1Q

https://disk.yandex.ru/d/oWlOciWiOoxOpQ

### Competing Interest Statement

The authors have declared no competing interest.

